# Nlrp6-dependent sentinel goblet cell functions dictate defensive responses during bacterial invasion of colonic crypts

**DOI:** 10.64898/2026.11.13.766363

**Authors:** Mahadevan Venkita Subramani, Gillian Blancke, Lars Vereecke, Jenny Gustafsson, Elisabeth E.L. Nyström, George M.H. Birchenough

## Abstract

The intestinal mucus layer serves as the primary physical barrier against microbial invasion, shielding the epithelium from luminal pathogens and inflammatory insults. This barrier is maintained by a heterogeneous population of goblet cells (GCs), which secrete regionally and organizationally distinct mucus layers. However, when this primary barrier is compromised, the burden of defense likely shifts to secondary mechanisms that remain incompletely defined. Therefore, we aimed to investigate the proposed role of microbe-sensing sentinel goblet cells (senGCs) as a secondary defensive mechanism that becomes critical when the primary mucus barrier fails, focusing on their capacity to mount localized secretory responses and preserve crypt sterility under challenge conditions. Using a GC specific *Nlrp6* knockout model (*Nlrp6^ΔGC^*), we demonstrate that senGC deficiency leads to increased susceptibility to DSS induced colitis and *Citrobacter rodentium* infection, marked by exacerbated inflammation and enhanced crypt invasion respectively. Transcriptomic and immunological profiling revealed both common and sex-specific immune dysregulation, with especially female *Nlrp6^ΔGC^*mice exhibiting heightened mucosal pathogen burden, reduced Th17 expansion and increased CD8⁺ T cell recruitment. The senGC-deficient mice also showed diminished baseline mucus secretion and accelerated GC loss during infection, suggesting a previously unrecognized mucus maintenance role for senGCs. Furthermore, infection induced expansion of Nlrp6⁺ GCs in wild-type mice, enhancing MAMP-triggered mucus secretion, a response that is absent in *Nlrp6^ΔGC^* tissue. These findings confirm senGCs as an important regulator of epithelial protection by integrating GC-intrinsic microbial sensing with mucosal defense.

## Introduction

The gastrointestinal tract is a site of constant microbial challenge, and its epithelial surface must balance nutrient absorption with robust defense. Central to this protection is the mucus barrier, a dynamic, regionally specialized structure primarily composed of the polymeric mucin Muc2, secreted by intestinal goblet cells (GCs) [1]. Rather than serving as a static shield, this mucus system adapts to the distinct physiological demands of each intestinal segment. In the small intestine, where nutrient uptake is prioritized, the mucus remains loosely organized and is fortified by antimicrobial peptides (AMPs) secreted by enterocytes and Paneth cells [2–4]. These components collectively limit microbial encroachment while preserving absorptive function.

In contrast, the distal colon faces a higher microbial load and relies on a more stratified mucus architecture. Here, the inner mucus layer (IML) forms a dense, bacteria-excluding matrix, while the outer mucus layer (OML) accommodates commensal communities. The IML is continuously replenished by surface GCs and plays a pivotal role in maintaining epithelial integrity [3]. However, this barrier is not impervious. Environmental stressors, microbial dysbiosis, and genetic disruptions in mucus production can render the IML penetrable, exposing the epithelium to microbial invasion and inflammation [5–7].

Goblet cells themselves are functionally diverse and spatially compartmentalized [8]. Single-cell transcriptomic analyses have revealed distinct GC lineages including canonical and non-canonical populations, based on their transcriptional profiles. Among canonical GCs, intercrypt goblet cells (icGCs) are terminally differentiated and rely on the transcription factor Spdef for their maturation [9]. These cells produce intercrypt mucus, a specialized sub-component of the IML that enhances its impermeability [8]. In line with this, loss of Spdef impairs icGC function, leading to defective mucus architecture and increased vulnerability to colonic inflammation. Another critical mucus substructure is the crypt plume, secreted by crypt-resident GCs and stabilized through transglutaminase 3 (Tgm3) mediated crosslinking of Muc2 [10]. Together, the intercrypt and crypt plume mucus systems form a multilayered defense that is both spatially and biochemically tuned. Yet, when this primary barrier fails, the host must rely on rapid compensatory mechanisms to prevent microbial access to the crypt niche.

One such mechanism is proposed to involve sentinel goblet cells (senGCs), a specialized GC subset positioned near crypt openings [11]. These cells are thought to act as microbial sensors, detecting microbial-associated molecular patterns (MAMPs) and initiating emergency mucus secretion. This response is orchestrated through a signaling cascade involving Toll-like receptors (TLRs), reactive oxygen species generation via Duox2 and Nlrp6 inflammasome activation [12]. Upon stimulation, senGCs undergo calcium-dependent compound exocytosis of Muc2, effectively flushing bacteria from the crypt entrance. The activated senGCs are ultimately shed out of the epithelium [11].

While senGCs have been proposed as key players in barrier reinforcement, their functional role during active infection remains underexplored *in vivo*. Given the importance of mucus integrity in preventing infection and colitis, it is crucial to understand how senGCs respond and further shape downstream GC functions when the IML is compromised. Additionally, this gap in knowledge limits our understanding of how the epithelium recovers from barrier disruption and how pathogens exploit these vulnerabilities.

Therefore, in this study, we investigate the role of senGCs as a functional secondary defensive mechanism following IML failure. We evaluate how senGCs influence GC physiology, mucus architecture, and host responses during infection. To do this, we use *Citrobacter rodentium*, a murine attaching and effacing pathogen that breaches the IML to colonize the colonic epithelium and invade crypts [13]. Additionally, this pathogen modulates Muc2 secretion and GC dynamics making it an ideal model for studying mucus barrier disruption, senGC activation and epithelial colonization [14]. Our findings reveal that senGCs orchestrate a dynamic defense program involving inducible mucus secretion, acute transcriptional reprogramming, and immune modulation. We show a previously unrecognized role for senGCs in sustaining baseline mucus secretion during infection in addition to protecting against *C. rodentium* driven epithelial damage.

## Results

### Generation of functional sentinel goblet cell (senGC) knockout mice

Within the colon, the mucus barrier comprises a loose outer mucus layer (OML) layer colonized by commensals, overlying a dense inner mucus layer (IML) that normally excludes microbes and inert particles as demonstrated by *ex vivo* imaging of colonic mucosa overlaid with 1µm fluorescent beads that accumulate exclusively at the surface of the mucus layer (Fig. 1A). Bacteria visualised by fluorescent cell staining are normally found localised in a discrete outer layer, likely representing the OML, whereas the tissue-adjacent IML region is devoid of bacterial cells (Fig 1A). However, IML sterility is not absolute, as bacterial penetration of the IML and colonisation down to the epithelial surface can be observed in otherwise healthy mice (Fig 1B), as well as under various conditions and disease states that are known to disrupt IML function.

**Figure 1:**
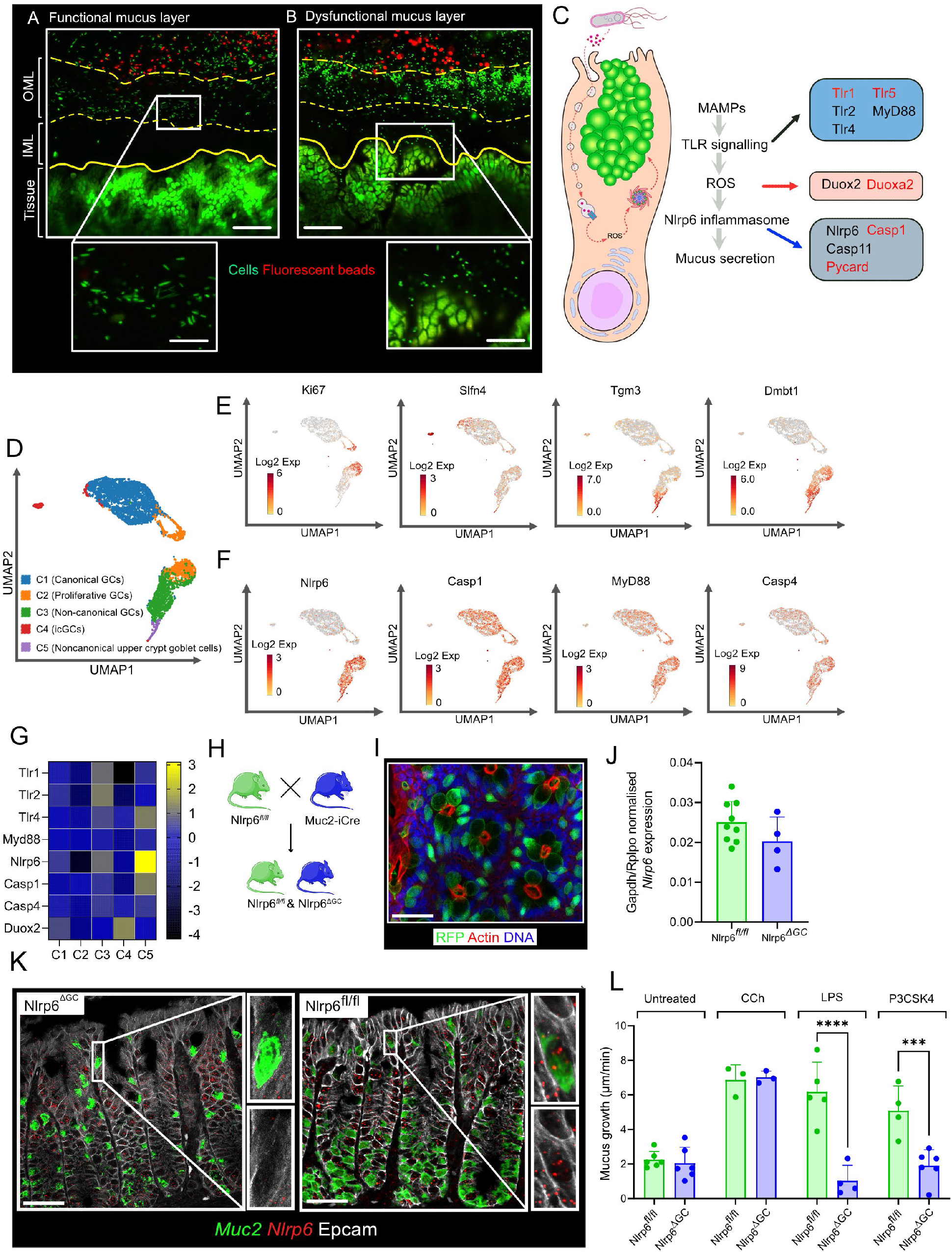
Generation and characterization of GC-specific Nlrp6 knockout mice. **A)** Representative confocal micrographs showing a functional colonic mucus barrier. Yellow lines indicate the boundaries of the outer mucus layer (OML) and inner mucus layer (IML). Enlarged regions are shown below. **B)** Representative confocal micrographs showing a dysfunctional colonic mucus barrier with fluorescent microbead penetration into the IML. Enlarged regions are shown below. **C)** Description of senGC activation pathway depicting known (black) and putative (red) pathway genes. **D)** Uniform Manifold Approximation and Projection (UMAP) of colonic GC transcriptomic data showing five GC sub-populations identified by clustering. **E)** UMAP plots showing the expression patterns of *Ki67*, *Slfn4*, *Tgm3* and *Dmbt1* across GC sub-populations. **F)** UMAP plots showing the expression patterns of senGC-associated genes *Nlrp6*, *Casp1*, *Myd88* and *Casp4* across GC populations. **G)** Heatmap showing normalized expression of selected senGC-associated genes across the five GC sub-populations (C1-C5). Colour scale indicates normalized gene expression (z-score). **H)** Schematic of the breeding strategy used to generate GC-specific Nlrp6 knockout mice (Nlrp6^ΔGC^) by crossing Nlrp6^fl/fl^ mice with Muc2-iCre mice. **I)** Representative confocal micrograph of distal colonic tissue from ROSA-tdRFP/Muc2-iCre reporter mice showing reporter expression in GCs. **J)** Relative *Nlrp6* mRNA expression quantified by qRT-PCR in colonic tissue from Nlrp6^fl/fl^ and Nlrp6^ΔGC^ mice, normalized to *Gapdh/Rplpo* expression. **K)** Representative mRNA in situ hybridization images of distal colonic tissue from Nlrp6^ΔGC^ and Nlrp6^fl/fl^ mice showing *Muc2* and *Nlrp6* expression. Enlarged regions are shown alongside the corresponding images. **L)** Ex vivo colonic mucus growth rate in Nlrp6^fl/fl^ and Nlrp6^ΔGC^ mice under untreated conditions or following stimulation with carbachol (CCh), lipopolysaccharide (LPS) or Pam3CSK4 (P3CSK4). Data represent n = 4-8 mice per group (J) or n = 4-6 mice per group per treatment (L). Data are pooled from two independent experiments (J). Bar graphs show mean ± SEM with individual data points overlaid (J, L). Statistical comparisons were performed using two-way ANOVA followed by Fisher’s LSD post hoc test (L). Statistical significance is indicated as p < 0.001 (***), p < 0.0001 (****). Image scale bars represent 20 µm (A, B, main images), 10 µm (A, B, insets), 40 µm (I) and 50 µm (K).

Sentinel goblet cells (senGC) have been proposed as a secondary GC intrinsic protective system that can orchestrate protective mucus secretion in response to bacterial IML penetration; however, this concept has yet to be fully validated *in* vivo due to lack of targeted senGC-deficient mouse models. We therefore set out to generate mice that lacked the secretory response reported in senGCs [11]. Activation of senGCs in response to MAMPs triggers downstream signaling events involving multiple mediators (Fig. 1C). As the expression of these genes is not restricted to goblet cells, a complete knockout would likely lead to stochastic senGC independent defects in the model system [12]. Moreover, GC specific knock out of the chosen mediator should ideally not compromise other functions in the heterogenous goblet cell population. We therefore reasoned that GC targeted conditional KO of one of these mediators would be suitable for generating a senGC deficient mouse model.

In order to identify a suitable candidate for conditional knockout, we mined goblet cell specific expression of senGC activation pathway genes from a recently published single cell transcriptomic study that identified both canonical (enriched in typical GC gene expression) and non-canonical (enriched in non-typical GC gene expression) GC differentiation trajectories [8]. Upon applying UMAP dimensionality reduction and K-means clustering algorithm, goblet cells were labelled as 5 distinct clusters (Fig. 1D). Cluster identity and crypt base-surface axis position was established by comparing gene expression of *Ki67, Slfn4, Tgm3, Dmbt1* (Fig. 1E, F). Cluster C2 had an enriched expression of proliferative stem cell marker *Ki67* classifying them as a group of deep-crypt proliferative GCs. We termed cluster C4 as icGCs based on their expression of *Slfn4 –* a marker for terminal differentiated surface GCs [8]. Clusters C3 and C5 closely associated with non-canonical GCs based on their increased expression of enterocyte linked gene *Dmbt1* [8]; Additionally, association of *Tgm3* mRNA with cluster C5 revealed them as upper crypt GCs. We then investigated markers that are involved in the senGC secretory pathway (*Tlr1, Tlr2, Tlr4, Nlrp6, MyD88, Casp1, Casp4, Duox2)* and detected variable expression across both canonical and non-canonical GC clusters (Fig. 1F, G). Notably, *Nlrp6* expression was specifically associated with the non-canonical clusters C3 and C5, and thus we determined that the upper crypt C5 cluster was most likely to include senGCs based on their position and expression of senGC activation mediators.

As Nlrp6 is a critical mediator of the senGC activation pathway, and shows restricted expression to non-canonical GCs, we reasoned that knockout of *Nlrp6* in GCs would provide a targeted senGC deficiency model. We therefore generated GC specific *Nlrp6* knock outs by crossing *Nlrp6^fl/fl^* with Muc2-iCre mice to produce *Nlrp6^ΔGC^* (Fig. 1H). We validated the selectivity of the Muc2-iCre strain examining the offspring of Muc2-iCre mice crossed with the Cre reporter strain ROSAtdRFP. In this strain, Cre selectively induces RFP signal in the cytosol of *Muc2+* GCs. Examination of whole mount colonic tissue from these mice identified restricted RFP expression in a subset of epithelial cells with distinct GC morphology, thus confirming the GC specificity of Cre expression (Fig. 1I).

To validate the model, we compared *Nlrp6* RNA expression in the colonic tissue of *Nlrp6^fl/fl^* and *Nlrp6^ΔGC^* mice and found a (non-significant) trend towards decreased *Nlrp6* expression in *Nlrp6^ΔGC^* (Fig. 1J). As the majority of *Nlrp6* expression in colon has been reported previously to be in the non-GC epithelial population [16], we sought to confirm deletion of *Nlrp6* in GCs using mRNA *in situ* hybridisation. Using this approach, we detected *Nlrp6* expressing GCs in *Nlrp6^fl/fl^* colonic tissue sections in the upper crypt region where senGCs are localized, which were absent in *Nlrp6^ΔGC^* tissues (Fig. 1K).

To functionally validate the knock-out model, we used an *ex vivo* mucus measurement assay, as previously established [17]. We treated colonic tissues from *Nlrp6^fl/fl^*and *Nlrp6^ΔGC^* with carbachol (a cholinergic secretagogue) and senGC-activating bacterial MAMPs (LPS, P3CSK4) and measured mucus growth in response to these compounds. Both MAMPs induced mucus secretion in *Nlrp6^fl/fl^* tissue, but not in *Nlrp6^ΔGC^*tissue, whereas tissue from both genotypes secreted mucus in response to carbachol (Fig. 1L), indicating that senGC-dependent secretory responses were specifically disabled in *Nlrp6^ΔGC^* mice. Together, our results confirm that *Nlrp6^ΔGC^* mice lack GC-specific *Nlrp6* expression and have disabled senGC-specific secretory responses and therefore represent a viable model to evaluate the function of senGCs *in vivo*.

### Nlrp6^ΔGC^ mice are more susceptible to chemical and biological colitogenic agents

Having validated the GC-specific Nlrp6 knockout model as deficient in senGCs and their unique mucus secretory responses, we next examined the contribution of this subpopulation to secondary mucosal defense during chemically induced barrier disruption and infection. Firstly, to evaluate senGC-mediated defense in a chemically induced model of barrier compromise, we exposed both *Nlrp6^fl/fl^* and *Nlrp6^ΔGC^* mice to dextran sodium sulfate (DSS), a colonic mucus barrier disruptor, for eight days and measured weight loss as a response to the treatment in addition to procuring tissue samples for calculating disease activity index (Fig. 2A).

**Figure 2:**
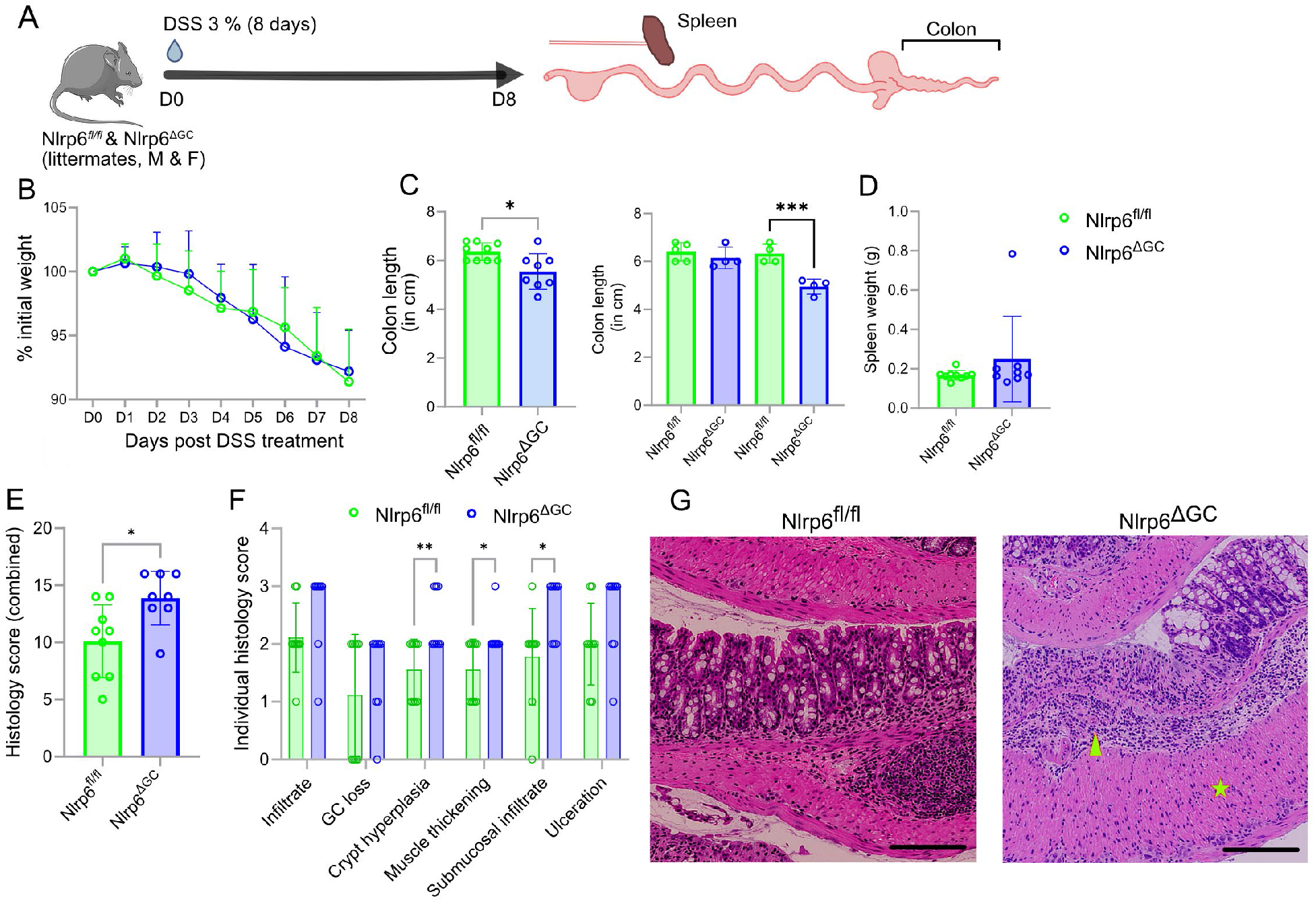
Nlrp6^ΔGC^ mice exhibit increased susceptibility to DSS-induced colitis. A) Experimental timeline showing administration of 3% dextran sulfate sodium (DSS) for 8 days in Nlrp6^fl/fl^ and Nlrp6^ΔGC^ mice, followed by sample collection. B) Daily body weight changes expressed as a percentage of initial body weight (mean ± SEM). C) Colon length at day 8 (mean ± SEM). Left, pooled genotype comparison by Mann-Whitney U test; right, sex-stratified comparison by one-way ANOVA followed by Fisher’s LSD post hoc test. D) Spleen weight at day 8 (mean ± SEM). E) Combined histopathology score of colonic damage (mean ± SEM), analysed using the Mann-Whitney U test. F) Individual histopathology scores for inflammatory infiltrate, GC loss, crypt hyperplasia, muscle thickening, submucosal infiltrate and ulceration (mean ± SEM), analysed using two-way ANOVA followed by Fisher’s LSD post hoc test. G) Representative H&E-stained distal colonic sections from Nlrp6^fl/fl^ and Nlrp6^ΔGC^ mice. Yellow asterisk indicates muscle thickening, and the arrowhead indicates submucosal inflammatory infiltration. Data represent n = 8-9 mice per genotype, including males and females from different litters. Data are presented as mean ± SEM, with individual data points overlaid where indicated. Statistical significance is indicated as p < 0.05 (*) and p < 0.01 (**). Image scale bars represent 100 µm (G).

While weight loss was similar between groups, *Nlrp6^ΔGC^*mice presented with significantly reduced colon lengths, a surrogate marker of increased tissue inflammation which drives colitis (Fig. 2B, C). Of note, female *Nlrp6^ΔGC^* seemed to be the most affected group when sex specific differences were compared (Fig. 2C). While spleen weights were unaffected, histological scoring revealed a more pronounced sex independent epithelial pathology in senGC-deficient mice, including exaggerated crypt hyperplasia, muscularis propria thickening, and heightened submucosal immune infiltration (Fig. 2D, E, F & G). Overall, these results show that *Nlrp6^ΔGC^* presented a moderately increased susceptibility to chemically induced barrier disruption, with indications of increased effects with female *Nlrp6^ΔGC^* mice.

Next, we evaluated senGC mediated defence against a biological colitogenic agent. To this end, we infected both *Nlrp6^fl/fl^* and *Nlrp6^ΔGC^* mice with *C. rodentium* over a 10-day regime, followed by assessment of stool and distal colonic tissue for bacterial quantification (Fig. 3A). As *C. rodentium* is an enteropathogen capable of breaching the mucus barrier and colonizing the colonic epithelium we expected that this pathogen would be ideally suited to activate and be modulated by the senGC response (Fig 3B).

**Figure 3:**
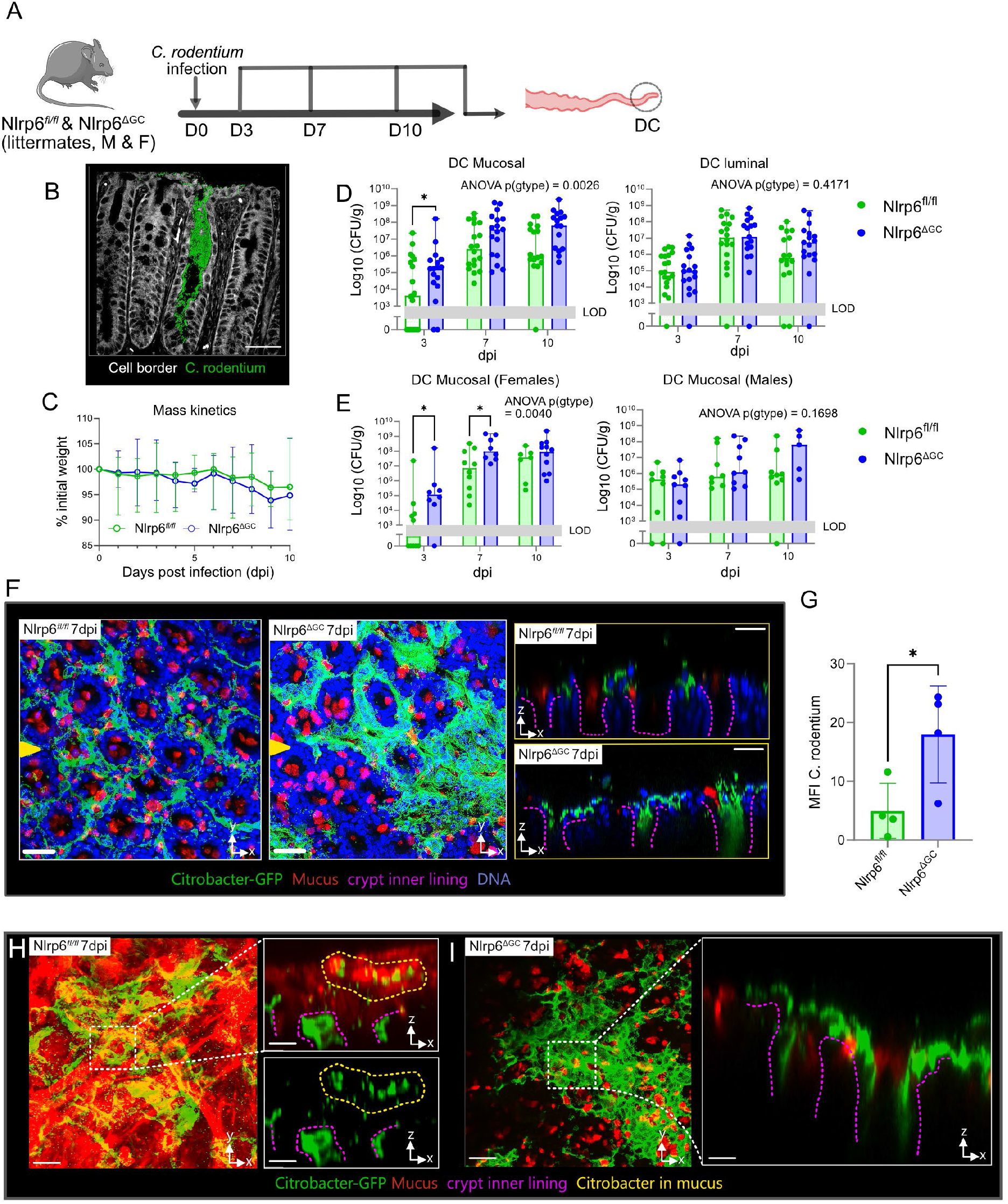
GC-specific Nlrp6 deficiency increases susceptibility to *C. rodentium* infection. A) Schematic of the experimental design showing *C. rodentium* infection of Nlrp6^fl/fl^ and Nlrp6^ΔGC^ mice, with distal colon tissue and stool collection at 3, 7 and 10 days post-infection (dpi). B) Representative confocal micrograph of distal colonic crypts showing GFP-expressing *C. rodentium* and EpCAM staining. C) Body weight changes in Nlrp6^fl/fl^ and Nlrp6^ΔGC^ mice following *C. rodentium* infection, expressed as a percentage of initial body weight. D) Mucosal (left) and luminal (right) *C. rodentium* burden quantified as colony-forming units per gram (CFU/g) in distal colon tissue and stool, respectively, at 3, 7 and 10 dpi. E) Sex-stratified analysis of distal colonic mucosal *C. rodentium* burden in female (left) and male (right) mice at 3, 7 and 10 dpi. F) Representative ex vivo confocal micrographs of distal colonic tissue from Nlrp6^fl/fl^ and Nlrp6^ΔGC^ mice at 7 dpi. Images show x/y-axis views (left) and x/z-axis cross-sections (right), illustrating GFP-expressing *C. rodentium* localization within colonic crypts. Yellow arrowheads indicate the positions of optical sections, and pink dashed lines outline crypt lumens. G) Quantification of GFP-expressing *C. rodentium* mean fluorescence intensity (MFI) in distal colonic tissue from Nlrp6^fl/fl^ and Nlrp6^ΔGC^ mice at 7 dpi. H) Representative ex vivo confocal micrographs of distal colonic tissue from Nlrp6^fl/fl^ mice at 7 dpi, showing *C. rodentium* localization within mucus and colonic crypts. Enlarged x/z-axis cross-sections are shown alongside, with yellow dashed lines indicating mucus-associated bacteria and pink dashed lines outlining crypt lumens. I) Representative ex vivo confocal micrographs of distal colonic tissue from Nlrp6^ΔGC^ mice at 7 dpi, showing *C. rodentium* localization within mucus and colonic crypts. Enlarged x/z-axis cross-sections are shown alongside, with pink dashed lines outlining crypt lumens. Data represent n = 9-13 mice per group across three independent infection experiments (C-E), n = 2 mice per group (F), or n = 4 mice per group (H, I). Data are presented as median and interquartile range (C-E) or mean ± SEM with individual data points overlaid (G). Statistical comparisons were performed using two-way ANOVA followed by Fisher’s LSD multiple comparisons test (C-E) or the Mann-Whitney U test (G). Statistical significance is indicated as p < 0.05 (\**), p < 0.01 (**), and p < 0.001 (\*\*\**). Grey shaded areas indicate the limit of detection (LOD). Image scale bars represent 30 µm (B), 100 µm (F, H, I; x/y-axis views) and 25 µm (F, H, I; x/z-axis cross-sections).

Across genotypes, weight loss post infection did not show any significant differences (Fig. 3C). 2-way ANOVA comparing genotype and days post-infection found that genotype significantly contributed to variation in mucosal bacterial load across the dataset, independent of time point, with *Nlrp6^ΔGC^* animals showing significantly higher mucosal bacterial burden compared to *Nlrp6^fl/fl^* animals at 3 dpi and a tendency towards higher burdens at both 7 and 10 dpi. In contrast, stool bacterial loads remained comparable across genotypes and time points, indicating that the observed mucosal differences are not reflected in luminal shedding (Fig. 3D). Given the well-established sex-dependent variation in murine responses to gastrointestinal disease, we next investigated whether *Nlrp6^ΔGC^* displayed differential susceptibility based on sex. Upon stratification, we observed that female *Nlrp6^ΔGC^* animals exhibited significantly higher mucosal bacterial loads at 3- and 7-days post-infection compared to *Nlrp6^fl/fl^* controls. In contrast, male *Nlrp6^ΔGC^* mice demonstrated bacterial burdens similar to *Nlrp6^fl/fl^* controls across all time points, suggesting sex-specific differences in *C. rodentium* colonisation in the absence of functional senGCs (Fig. 3E).

To investigate whether this increased bacterial load in *Nlrp6^ΔGC^*mice associated with crypt-localized invasion, we infected *Nlrp6^fl/fl^*and *Nlrp6^ΔGC^* mice with GFP-expressing *C. rodentium* and examined distal colonic whole-mounts at 7 dpi using confocal microscopy. Tissue was stained for mucus (UEA1) and nuclei (Hoechst), allowing visualization of bacterial distribution relative to the epithelial architecture. *Nlrp6^fl/fl^*animals showed bacterial accumulation primarily restricted to intercrypt regions, with crypt entrances remaining largely uncolonized. However, female *Nlrp6^ΔGC^* mice displayed prominent bacterial infiltration within crypt openings (Fig. 3F). Quantitative analysis of bacterial signal intensity revealed significantly higher *C. rodentium* tissue association in *Nlrp6^ΔGC^ animals*, correlating the enhanced crypt invasion (Fig. 3G).

We next sought to determine whether the absence of Nlrp6-dependent senGC mucus secretion contributed directly to this phenotype. To this end, we performed *ex vivo* lectin staining of distal colonic tissues from infected WT and senGC-deficient female mice. In *Nlrp6^fl/fl^* samples, we observed robust co-localization of bacteria and mucus at crypt entrances, indicative of a senGC-mediated compound secretory response (Fig. 3H). Conversely, in *Nlrp6^ΔGC^* tissue, such co-localization was absent, and bacterial infiltration into crypts was more frequent. These observations suggest that senGC-driven mucus secretion plays a critical role in restricting bacterial access to specific epithelial niches during acute infection (Fig. 3I).

Taken together, these results demonstrate that senGCs play a central and non-redundant role in maintaining crypt barrier integrity during both chemically induced mucosal challenge and microbial invasion, with their absence leading to enhanced crypt invasion, impaired pathogen clearance, and exacerbated inflammatory pathology, with enhanced pathogen burdens particularly noted in female mice. As *C. rodentium* infection represents a more biologically relevant model than DSS for investigating mucosal defence, we selected this model for further investigation.

### senGC deficiency drives sex-specific baseline and infection driven transcriptomic alterations

Since we observed a sex-dependent sensitivity to *C. rodentium* infection among *Nlrp6^ΔGC^* animals, we next investigated basal transcriptomic differences between female *Nlrp6^ΔGC^* and female *Nlrp6^fl/fl^* mice in addition to a parallel comparison in males. In uninfected samples, principal component analysis (PCA) separated mice of both genotypes based primarily on sex indicating overall distinct baseline transcriptional profiles between the sexes (Fig. 4A). Volcano plot analysis of female *Nlrp6^ΔGC^* (n=2) versus *Nlrp6^fl/fl^* (n=3) samples identified multiple significantly upregulated genes, including several immunoglobulin genes (*Igha*, *Igkv8-28*, *Igkv7-33*) alongside goblet cell associated transcripts *Reg4, Zg16 and Ang4* (Fig. 4B, left). In contrast, male *Nlrp6^ΔGC^* (n=4) versus *Nlrp6^fl/fl^* (n=4) comparisons yielded only a single differentially expressed gene (*Slc4a1*), underscoring a female-specific baseline alteration in immune and goblet cell gene networks (Fig. 4B, right). Gene ontology (GO) enrichment analysis of genes upregulated in female *Nlrp6^ΔGC^* mice revealed significant overrepresentation of immune-related processes, including immune system process, B cell– mediated adaptive immunity, lymphocyte-mediated immunity, and response to stimulus (Fig. 4C).

**Figure 4:**
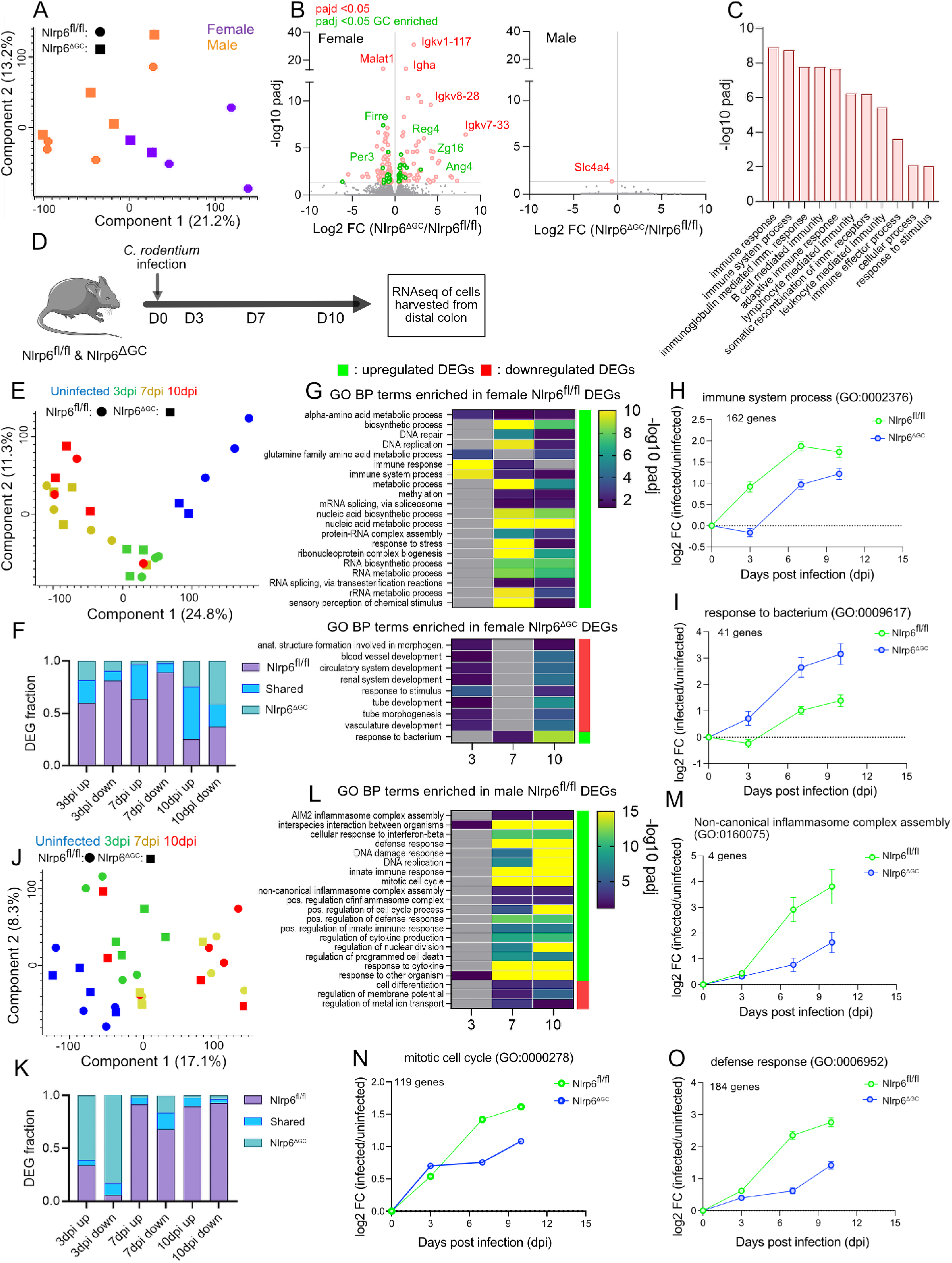
GC-specific Nlrp6 deficiency drives sex-specific baseline and infection-induced transcriptomic alterations. A) Principal component analysis (PCA) of RNA-seq data from distal colonic cells of uninfected female and male Nlrp6^fl/fl^ and Nlrp6^ΔGC^ mice. Samples are colour-coded by sex and shaped by genotype. B) Volcano plots showing differential gene expression between uninfected Nlrp6^ΔGC^ and Nlrp6^fl/fl^ mice in females (left) and males (right). The x-axis indicates log₂ fold change (log₂FC), and the y-axis represents -log₁₀ adjusted p-value (padj). Differentially expressed genes (padj < 0.05) are highlighted in red, with GC-enriched genes indicated in green. C) Gene Ontology (GO) biological process enrichment analysis of differentially expressed genes in uninfected female Nlrp6^ΔGC^ mice compared with Nlrp6^fl/fl^ controls. Bars indicate -log₁₀ padj. D) Schematic of the experimental design showing *C. rodentium* infection and distal colonic tissue collection at 3, 7 and 10 days post-infection (dpi) for RNA sequencing. E) PCA of distal colonic transcriptomic data from female Nlrp6^fl/fl^ and Nlrp6^ΔGC^ mice at uninfected, 3, 7 and 10 dpi. F) Proportions of differentially expressed genes (DEGs; padj < 0.05) unique to Nlrp6^fl/fl^, shared between genotypes, or unique to Nlrp6^ΔGC^ female mice at 3, 7 and 10 dpi. G) Heatmap showing GO biological process enrichment among upregulated and downregulated DEGs in female mice across infection time points. Colour scale indicates -log₁₀ padj. H-I) Mean log₂FC of genes associated with immune system process (H; GO:0002376), response to bacterium (I; GO:0009617) in female mice during infection. J) PCA of distal colonic transcriptomic data from male Nlrp6^fl/fl^ and Nlrp6^ΔGC^ mice at uninfected, 3, 7 and 10 dpi. K) Proportions of DEGs unique to Nlrp6^fl/fl^, shared between genotypes, or unique to Nlrp6^ΔGC^ male mice at 3, 7 and 10 dpi. L) Heatmap showing GO biological process enrichment among upregulated and downregulated DEGs in male mice across infection time points. Colour scale indicates -log₁₀ padj. M–O) Mean log₂FC of genes associated with non-canonical inflammasome complex assembly (M; GO:0160075), mitotic cell cycle (N; GO:0000278) and defense response (O; GO:0006952) in male mice during infection. Data represent n = 2-4 mice per sex and genotype (A-C). Each point represents one animal in the PCA plots (A, E, J). Gene expression changes are expressed as log₂FC relative to the corresponding uninfected genotype (H, I, M-O), with lines showing mean values and error bars where indicated. Differential expression was assessed using adjusted p-values (padj < 0.05).

To characterize the infection response, *Nlrp6^fl/fl^* and *Nlrp6^ΔGC^* mice were orally infected with *Citrobacter rodentium* and distal colonic epithelial cells harvested at uninfected, 3-, 7-, and 10-days post-infection (dpi) for bulk RNA sequencing (Fig. 4D). Firstly, in female samples, PCA separated uninfected from infected time points along component 1, with 3 dpi profiles occupying an intermediate position and 7–10 dpi samples shifting further along the infection axis (Fig. 4E). To account for altered gene expression at baseline between genotypes, we focused our analysis on comparison of differentially expressed genes (DEGs) between infected mice and their genotype-specific uninfected controls. We then quantified the fraction of DEGs unique to *Nlrp6^fl/fl^*, shared between genotypes, or unique to *Nlrp6^ΔGC^* at each time point. *Nlrp6^fl/fl^*-specific DEGs dominated both upregulated and downregulated categories across 3 and 7 dpi, whereas *Nlrp6^ΔGC^*samples exhibited markedly fewer unique DEGs, perhaps suggesting a less complex infection-driven transcriptional program in *Nlrp6^ΔGC^* mice (Fig. 4F).

To functionally annotate the DEGs enriched in female *Nlrp6^fl/fl^* and female *Nlrp6^ΔGC^* animals across the infection we performed functional enrichment analysis for biological processes. Gene ontology (GO) analysis of upregulated DEGs in female *Nlrp6^fl/fl^* samples showed strong enrichment for genes associated with immune system process throughout the infection (Fig. 4G). On the contrary, upregulated DEGs in female *Nlrp6^ΔGC^* animals showed strong upregulation of response to bacterium genes (Fig. 4G). To illustrate the kinetics of selected biological processes, we plotted the mean log₂ fold change of all DEGs annotated to “immune system process” (162 genes) and “response to bacterium” (41 genes) over the infection time course in both genotypes. DEGs unique to female *Nlrp6^fl/fl^* mice revealed accelerated infection dependent enrichment of immune system-related gene expression, suggesting a more rapid immune response driven by senGC activation (Fig. 4H). In contrast, DEGs unique to female *Nlrp6^ΔGC^* mice showed an infection induced enrichment for bacterial response pathways, which may reflect a compensatory transcriptional program triggered by increased pathogen burden in the absence of senGC protection (Fig. 4I).

We next examined whether male *Nlrp6^fl/fl^* and *Nlrp6^ΔGC^* mice responded similarly to *C. rodentium* infection. PCA demonstrated that uninfected *Nlrp6^fl/fl^* and *Nlrp6^ΔGC^* samples clustered together, while infected samples diverged progressively in line with increasing days post-infection (Fig. 4J). In males, *Nlrp6^ΔGC^* -specific DEGs dominated at 3 dpi (although with no significant biological process associations), whereas *Nlrp6^fl/fl^*-specific DEGs dominated at 7 and 10 dpi (Fig. 4K). Examination of selected biological processes over-represented in *Nlrp6^fl/fl^*unique DEGs indicated enrichment of genes associated with multiple biological processes, including “non-canonical inflammasome complex assembly”, “defense response”, and “mitotic cell cycle processes” (Fig. 4L). All three biological processes were induced more rapidly and to a greater extent in male *Nlrp6^fl/fl^* mice than in *Nlrp6^ΔGC^* mice (Fig. 4M-O).

Overall, our data indicates that loss of senGCs drives a female-specific baseline shift in colonic tissue gene expression marked by upregulation of immunoglobulin and goblet cell-enriched transcripts. Furthermore, upon *C. rodentium* infection, both sexes exhibit a genotype-dependent differential transcriptional program, thus indicating that senGC function regulates tissue responses to infection with a sex-dependent manner.

### Mice deficient in senGCs possess altered baseline neutrophil levels and infection induced T cell dynamics

Our bulk RNA-seq analysis indicated altered immune processes in both baseline and infected senGC deficient mice. Moreover, since Nlrp6 is involved in inflammasome assembly and the maturation of inflammatory cytokines, we hypothesized that its deletion in goblet cells could impair local immune recruitment and epithelial defense during C. rodentium infection. To test this, we isolated lamina propria from the distal colon of *Nlrp6^fl/fl^* and *Nlrp6^ΔGC^* mice at steady state and seven days post-infection (7 dpi) and profiled their immune cell composition by flow cytometry (Fig. 5A).

**Figure 5:**
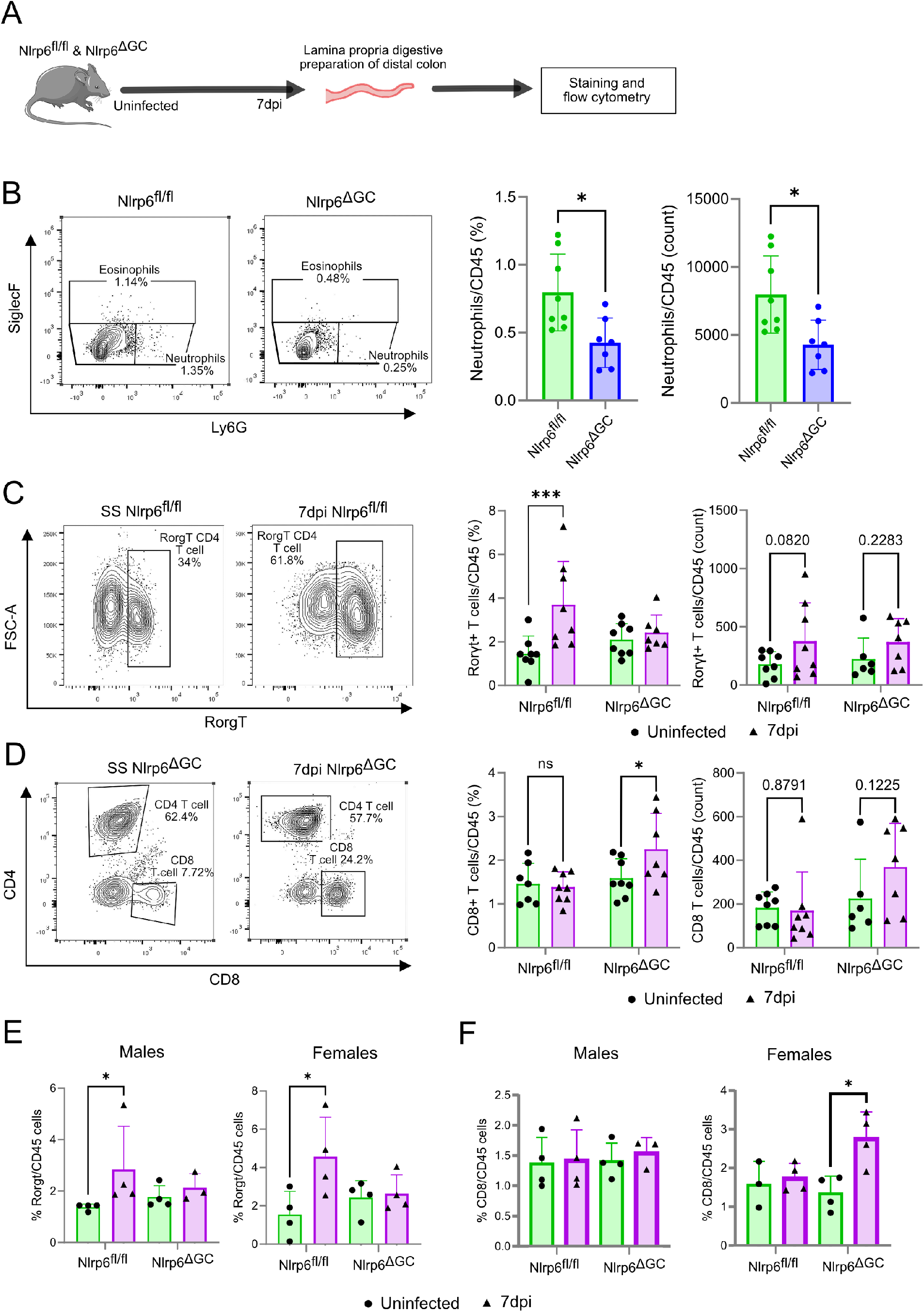
GC-specific Nlrp6 deficiency alters baseline neutrophil levels and infection-induced T cell dynamics. A) Experimental workflow for distal colonic lamina propria digestion, staining and flow cytometry analysis. B) Neutrophils (Ly6G⁺SiglecF⁻) among CD45⁺ cells in uninfected Nlrp6^fl/fl^ and Nlrp6^ΔGC^ mice. Representative flow plots (left) and quantification of frequency and absolute counts (right). C) RORγt⁺CD4⁺ T cells among lamina propria leukocytes of Nlrp6^fl/fl^ and Nlrp6^ΔGC^ mice. Representative flow plots (left) and relative and absolute cell quantification (right). D) CD8⁺ T cells among CD45⁺ cells. Representative gating plots (left) and relative and absolute CD8⁺ T cell quantification (right). E) Sex-stratified RORγt⁺ T cell frequencies in male and female Nlrp6^fl/fl^ and Nlrp6^ΔGC^ mice at steady state and 7 dpi. F) Sex-stratified CD8⁺ T cell frequencies in male and female Nlrp6^fl/fl^ and Nlrp6^ΔGC^ mice at steady state and 7 dpi. Data represent n = 7-8 mice per group per condition. Bars show mean ± SD. Statistical comparisons were performed using the Mann-Whitney U test; p < 0.05 (*), p < 0.001 (***); ns, not significant.

We employed a broad myeloid and B cell panel including eosinophils, neutrophils, monocytes, macrophages, dendritic cells, B cells, and CD138+ plasma cells, and a parallel T cell panel (CD3, CD4, CD8, Rorγt, Foxp3, T-bet, GATA3). At steady state, *Nlrp6^ΔGC^* mice displayed a significant reduction in both the percentage and absolute number of neutrophils among CD45+ cells (Fig. 5B). However, no other myeloid or B cell populations differed by genotype, nor did we observe any sex dependent differences in those populations (data not shown).

When we examined the T cell compartment, *Nlrp6^fl/fl^* mice mounted a robust Th17 response to *C. rodentium* marked by a significant increase in *Rorγt*+ CD4+ T cells, compared to uninfected *Nlrp6^fl/fl^*animals. This Th17 expansion was absent in *Nlrp6^ΔGC^* mice, indicating a failure to elicit this protective arm of T cell immunity during *C. rodentium* infection (Fig. 5C). Additionally, upon infection *Nlrp6^ΔGC^* mice exhibited a significantly higher proportion of cytotoxic CD8+ T cells compared to uninfected *Nlrp6^ΔGC^*animals, that was absent in *Nlrp6^fl/fl^* animals (Fig. 5D). Next, we stratified the data to investigate sex-dependent differences in our results. We observed that both male and female *Nlrp6^fl/fl^* mice displayed a significant increase in Rorγt⁺ CD45⁺ cells at 7 dpi compared to uninfected *Nlrp6^fl/fl^* controls (Fig. 5E). In the CD8 T cell compartment, female *Nlrp6^ΔGC^*mice exhibited a significant rise in CD8⁺ T cell frequencies at 7 dpi compared to uninfected female *Nlrp6^ΔGC^* mice, with male mice remaining unchanged compared to their controls (Fig. 5F).

Overall, these results demonstrate that senGC deficiency disrupts steady-state neutrophil homeostasis, blunts the *C. rodentium*-induced protective expansion of Th17 cells, and drives a compensatory rise in CD8⁺ T cells during infection in females. Together, these findings demonstrate that senGCs regulate local mucosal immune functions during *C. rodentium* infection.

### Maintenance of goblet cell frequency and mucus barrier function is regulated by senGCs during infection

Our results revealed that *C. rodentium* infection induces differential tissue responses in *Nlrp6^ΔGC^* animals. Tissue responses to *C. rodentium* infection normally include a variety of regiospecific responses such as epithelial crypt hyperplasia and mucus secretion aimed at protection of the host and elimination of the pathogen [18], [19]. Therefore, we examined if gross tissue responses and GC dynamics and mucus barrier properties varied between *Nlrp6^fl/fl^*and *Nlrp6^ΔGC^* animals.

To test the influence of active senGCs on colonic tissue responses to infection, we generated swiss-roll sections from *Nlrp6^fl/fl^*and *Nlrp6^ΔGC^* mice at 0 (uninfected), 3, 7, and 10 dpi and performed Alcian Blue/Periodic Acid– Schiff (AB/PAS) staining to localize mucin-filled goblet cells. Furthermore, we divided the tissue into proximal colon, proximal mid-colon, mid colon, and distal colon regions to look for regionalized differences in tissue parameters (Fig. 6A–B). We quantified crypt length and goblet cell frequency (number of mucin-filled goblet cells per unit crypt length). As expected, both genotypes showed infection induced crypt hyperplasia in the mid and distal colon, peaking at 7 dpi and partially regressing by 10 dpi, with no significant differences in magnitude or kinetics between *Nlrp6^fl/fl^*and *Nlrp6^ΔGC^* mice (Fig. 6C). In contrast, mucin filled goblet cell frequency declined significantly by 3 dpi across proximal, mid, and distal regions in *Nlrp6^ΔGC^* mice, whereas *Nlrp6^fl/fl^* animals maintained baseline goblet cell numbers until 7 dpi, when a modest decrease appeared restricted to the mid colon (Fig. 6D).

**Figure 6:**
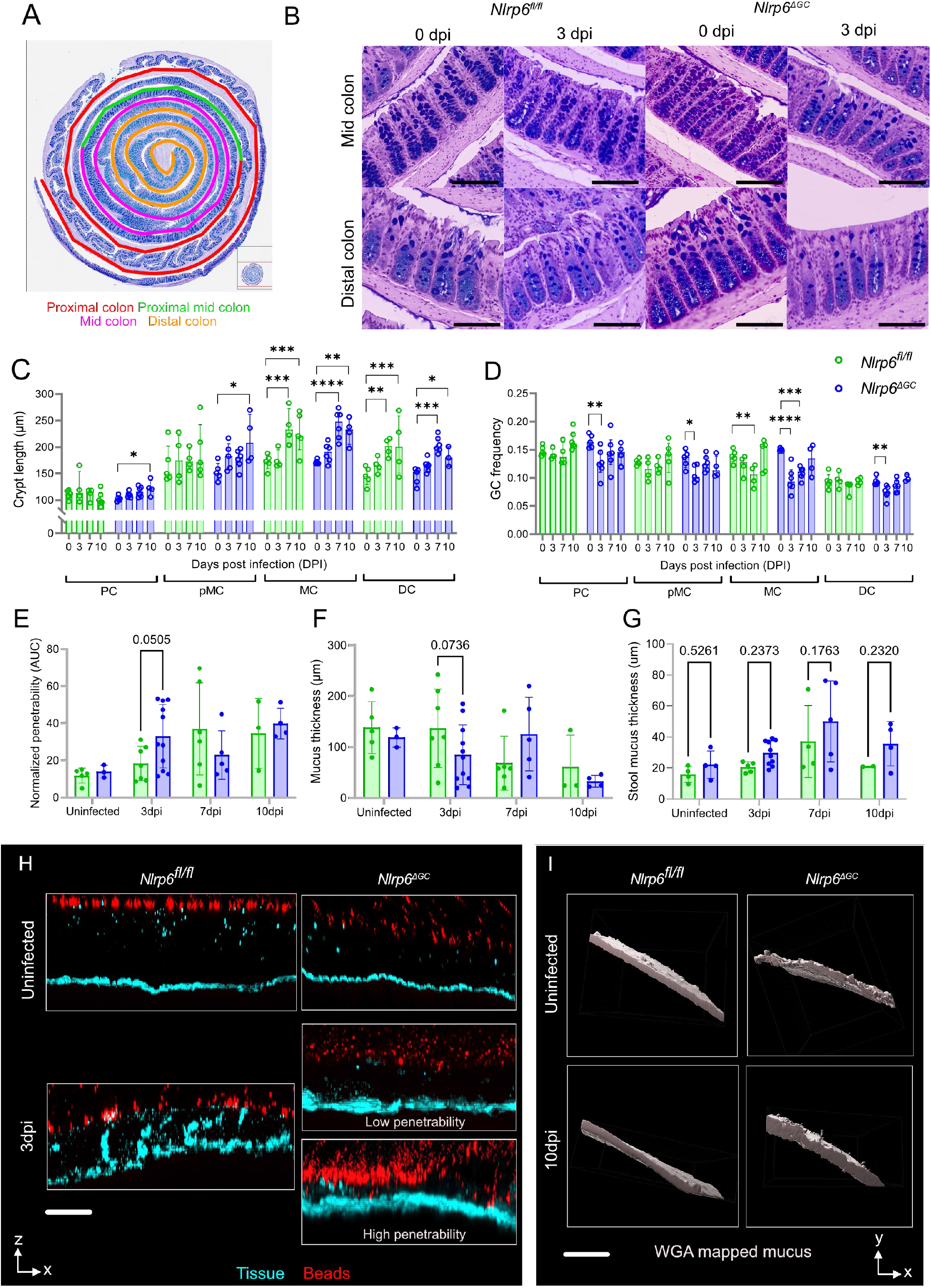
GC-specific Nlrp6 deficiency alters colonic crypt morphology and mucus barrier function during infection. A) Schematic of colonic regions analysed using Swiss rolls: proximal colon (PC), proximal mid colon (pMC), mid colon (MC) and distal colon (DC). B) Representative AB-PAS-stained sections of mid and distal colon from Nlrp6^fl/fl^ and Nlrp6^ΔGC^ mice at 0 and 3 days post-infection (dpi). C) Crypt length across different colonic regions at 0, 3, 7 and 10 dpi. Each data point represents the average of 15 crypts per region. D) GC frequency normalized to crypt length across different colonic regions at 0, 3, 7 and 10 dpi. Each data point represents the average of 10 crypts per region. E) IML penetrability quantified by ex vivo fluorescent microbead penetration, expressed as normalized area under the curve (AUC), at uninfected, 3, 7 and 10 dpi. F) IML thickness quantified by ex vivo mucus measurements at uninfected, 3, 7 and 10 dpi. G) Stool mucus thickness measured ex vivo at uninfected, 3, 7 and 10 dpi. H) Representative confocal x/z-axis cross-sections of distal colonic tissue from uninfected and infected Nlrp6^fl/fl^ and Nlrp6^ΔGC^ mice at 3 dpi, showing tissue (cyan) and fluorescent microbeads (red). Images illustrate low and high mucus penetrability in Nlrp6^ΔGC^ mice. I) Representative three-dimensional iso-surface reconstructions of WGA-labelled stool mucus from uninfected and infected Nlrp6^fl/fl^ and Nlrp6^ΔGC^ mice at 10 dpi. Data represent n = 4-6 mice per group per time point (C, D) and approximately n = 3-11 mice per group per time point (E-G). Data are presented as median and interquartile range (C, D) or mean ± SEM (E-G). Statistical comparisons were performed using two-way ANOVA followed by Fisher’s LSD multiple comparisons test (C, D) or the Mann–Whitney U test (E-G). Statistical significance is indicated as p < 0.05 (*), p < 0.01 (**), p < 0.001 (***) and p < 0.0001 (****). Exact p-values for non-significant comparisons are indicated where shown. Image scale bars represent 80 µm (B), 50 µm (H) and 100 µm (I).

To assess whether early goblet cell loss among *Nlrp6^ΔGC^*animals affected mucus barrier integrity, we performed ex vivo bead-penetrability assays on colonic tissues from both genotypes throughout the infection regime. Normalized bead penetrability increased, and mucus thickness decreased over the course of infection in *Nlrp6^fl/fl^* and *Nlrp6^ΔGC^* alike, reflecting progressive inner mucus layer (IML) barrier dysfunction (Fig. 6E, F, H). Interestingly, *Nlrp6^ΔGC^* tissues trended toward higher penetrability and lower thickness at 3 dpi, although this did not reach statistical significance with the current replicate number due to heterogeneity within the *Nlrp6^ΔGC^* group. Since tissue-level mucus alterations can influence the fecal mucus layer (FML) and modulate elimination kinetics of the pathogen, we measured FML thickness across infection. No significant genotype differences were observed at any time point; however, a non-significant upward trend in FML thickness emerged in *Nlrp6^ΔGC^* mice at 10 dpi, potentially indicating enhanced compensatory mucus secretion from regions proximal to distal colon (Fig. 6G & I).

Overall, these results demonstrate that *C. rodentium* infection induces genotype independent crypt hyperplasia in mid and distal colon; however, senGC deficiency was associated with reduction in mucin-filled GC frequency in mid and distal colon as early as 3-DPI and a tendency towards faster IML disruption during infection. Consequently, our data supports the concept that senGC function is essential for preserving goblet cell numbers early in infection and potentially delaying initial mucus barrier dysfunction induced by *C. rodentium*.

### Infection induces dynamic alterations in senGC location and function

Our results indicated that senGC function influences downstream tissue associated, goblet cell and IML responses to *C. rodentium* infection. As some of these responses may be regulated by dynamic changes in the senGC population itself, we lastly investigated how senGCs and senGC-dependent secretory responses were impacted by the infection.

To test this, we first performed mRNA in situ hybridization for *Nlrp6* and *Muc2* on colonic sections from uninfected and infected WT mice to localize and quantify senGCs in the epithelium (Fig. 7A). In uninfected tissue, *Nlrp6* transcripts localized predominantly to upper-crypt GCs with some signal in the crypt base GCs. However, by 3 dpi overall *Nlrp6* expression rose markedly, with a clear expansion of *Nlrp6⁺/Muc2⁺* GCs at the deep crypt regions. This elevated expression persisted at 7 and 10 dpi, indicating that *C. rodentium* infection induces expansion of *Nlrp6*-expressing GCs predominantly in the lower regions of the crypt (Fig. 7A).

**Figure 7:**
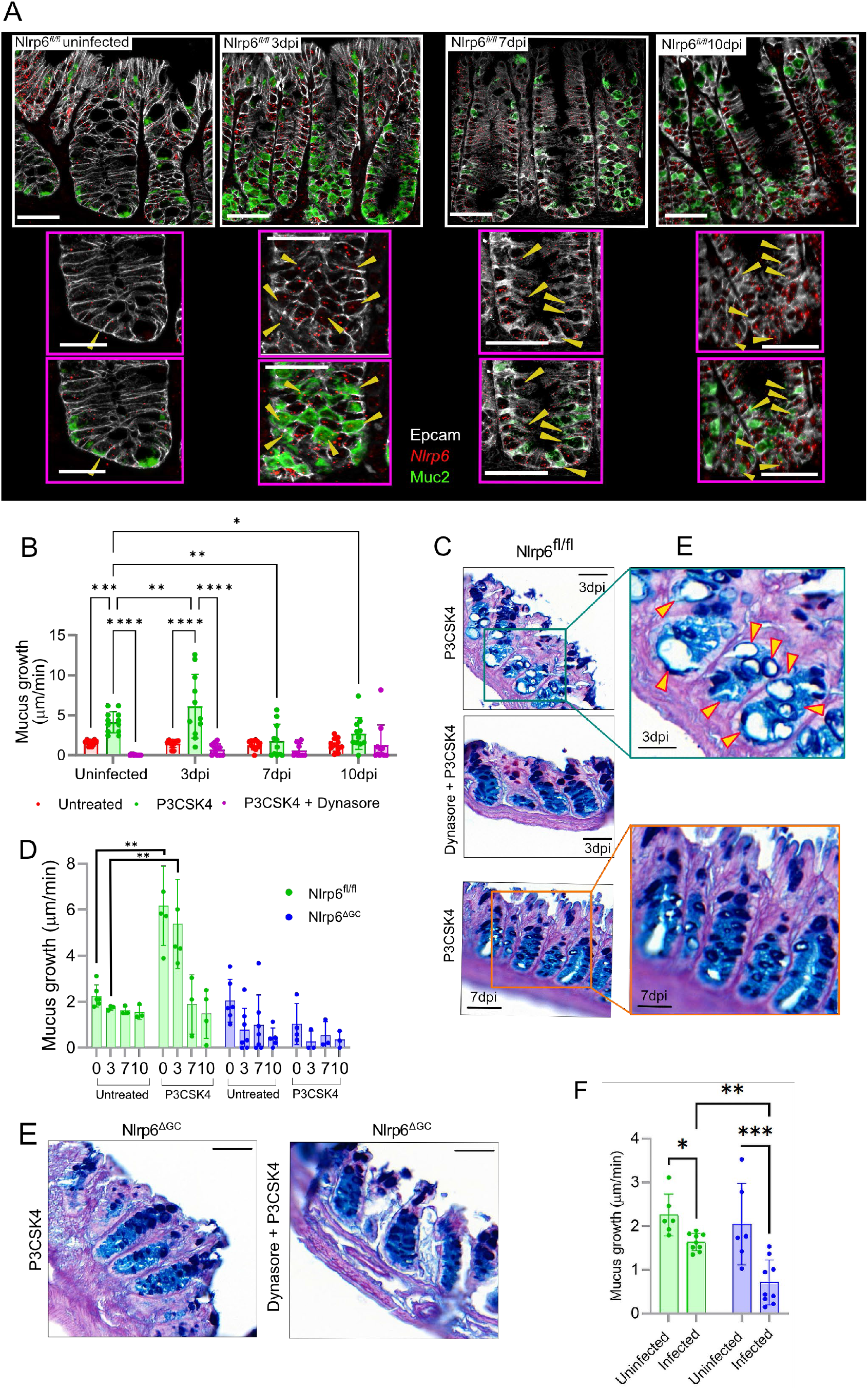
Infection induces dynamic alterations in senGC localization and function. A) mRNA in situ hybridization of Nlrp6 (red) and Muc2 (green) in colonic tissue from Nlrp6^fl/fl^ mice at uninfected, 3, 7 and 10 days post-infection (dpi), counterstained for EpCAM (white). Yellow arrowheads indicate Nlrp6⁺ GCs. Pink-bordered images show enlarged crypt regions. B) Ex vivo mucus growth in distal colonic tissue from Nlrp6^fl/fl^ mice at uninfected, 3, 7 and 10 dpi following treatment with the TLR2 agonist P3CSK4, alone or in combination with the endocytosis inhibitor Dynasore. C) Representative AB-PAS-stained colonic sections from Nlrp6^fl/fl^ mice at 3 and 7 dpi following ex vivo treatment with P3CSK4 with or without Dynasore. Enlarged regions highlight deep-crypt GC cavitation (yellow arrowheads), indicative of MAMP-induced mucus secretion. D) Ex vivo mucus growth in distal colonic tissue from Nlrp6^fl/fl^ and Nlrp6^ΔGC^ mice at uninfected, 3, 7 and 10 dpi, with or without P3CSK4 stimulation. E) Representative AB-PAS-stained colonic sections from Nlrp6^ΔGC^ mice at 3 dpi following ex vivo treatment with P3CSK4, alone or in combination with Dynasore, showing the absence of deep-crypt GC cavitation. F) Baseline ex vivo mucus growth in distal colonic tissue from Nlrp6^fl/fl^ and Nlrp6^ΔGC^ mice under uninfected conditions and following C. rodentium infection. Data represent n = 12-13 mice per group from two independent experiments (B), n = 5 mice per genotype (D), or n = 6-9 mice per group (F). Data are presented as mean ± SEM. Statistical comparisons were performed using two-way ANOVA followed by Fisher’s LSD post hoc test (B, D, F). Statistical significance is indicated as p < 0.05 (*), p < 0.01 (**), p < 0.001 (***) and p < 0.0001 (****). Image scale bars represent 40 µm (A, main images), 30 µm (A, enlarged regions) and 50 µm (C, E).

Next, to determine whether this shift in *Nlrp6⁺* GC frequency associated with functional changes in senGC-dependent mucus secretion, we examined colonic tissues from WT mice at 0, 3, 7, and 10 dpi and measured mucus growth *ex vivo* in response to the TLR2 agonist P3CSK4, with or without the endocytosis inhibitor Dynasore that blocks senGC-dependent secretory response to P3CSK4. Tissues from 3 dpi WT mice responded to P3CSK4 with a comparable mucus growth to uninfected controls, and this inducible secretion was completely abolished by Dynasore (Fig. 7B). Tissues from 7 and 10 dpi failed to show further enhancement over their respective baselines, suggesting a transient or exhaustion-prone secretory response during mid-infection (Fig. 7B).

To visualize the cellular origin of improved P3CSK4-induced secretion in WT animals during infection, we fixed and sectioned tissue treated with P3CSK4 (± Dynasore) and performed AB/PAS staining. In WT tissue at 3 and (to a lesser extent) at 7 dpi, P3CSK4 triggered cavitation in deep-crypt GCs indicative of secretion, a pattern absent in Dynasore-pretreated tissues (Fig. 7C), raising the possibility that MAMP-induced secretory response is altered at this stage of infection. Overall, these results indicate that the deep crypt *Nlrp6+* GCs induced by infection have a role to play in enhanced stimulated mucus secretion during early infection in WT animals.

Lastly, to confirm the role of the senGC activation pathway in the MAMP-induced mucus secretion during infection, we compared untreated and P3CSK4 treated tissues from *Nlrp6^fl/fl^*and *Nlrp6^ΔGC^* mice at each infection time point. As expected, *Nlrp6^fl/fl^* tissues treated with P3CSK4 showed a significant increase in mucus secretion compared to untreated tissue (Fig. 5D). *Nlrp6^ΔGC^*tissue did not respond to P3CSK4 stimulation at any stage of the infection, confirming the dependence of this inducible secretion on senGCs (Fig. 7D). Consistently, P3CSK4-treated *Nlrp6^ΔGC^* tissue lacked GC cavitation, confirming that senGCs are essential for initiating the deep crypt secretory response observed in 3 & 7dpi tissue treated with P3CSK4

(Fig. 7E). Surprisingly, tissue from *Nlrp6^ΔGC^* animals consistently showed significantly lower baseline mucus growth than *Nlrp6^fl/fl^* during infection (Fig. 7F). These results suggest that despite the loss of the MAMP-inducible secretory response during infection, senGCs may switch to maintaining baseline secretion and/or expansion of mucus, indicating that their specific functional role might be modulated in response to infection.

To summarize, these results demonstrate that *C. rodentium* infection drives an induction of *Nlrp6*-expressing GCs into the lower parts of crypt that participate in Nlrp6-dependent mucus secretion in response to microbial stimuli during early infection. This early, localized, and improved secretory response is lacking in *Nlrp6^ΔGC^* animals and may thus explain their increased vulnerability to barrier disruption. Furthermore, these findings indicate that that senGCs are essential for sustaining baseline mucus secretion during infection, unveiling a previously unrecognized role for this cell type.

## Discussion

In this study, we generated and validated a novel senGC deficiency model by goblet cell–specific knockout of *Nlrp6* (*Nlrp6^ΔGC^*). We subsequently used this model to demonstrate that senGCs are essential for mounting a rapid, localized secretory response to microbial stimuli in vivo. Their absence results in heightened susceptibility to both chemically induced colitis and *Citrobacter rodentium* infection. The senGC-deficient mice exhibited increased mucosal burden, impaired crypt sterility, and exacerbated inflammatory pathology. Bulk RNA sequencing revealed sex-specific baseline shifts and attenuated infection-induced immune responses in *Nlrp6ΔGC* mice, including reduced Th17 expansion and compensatory recruitment of cytotoxic CD8⁺ T cells. Additionally, senGC deficiency disrupted neutrophil homeostasis and accelerated goblet cell loss and inner mucus layer deterioration during infection. Notably, *C. rodentium* infection induced a dynamic expansion of *Nlrp6⁺* GC into the deep crypts of wild-type mice, enhancing MAMP-triggered mucus secretion, a response entirely absent in *Nlrp6^ΔGC^* tissue.

A major finding is that female *Nlrp6^ΔGC^* mice exhibit more severe colitis and greater susceptibility to *C. rodentium* infection than their male counterparts, as reflected in microbial loads and histological scores. Flow cytometric profiling of the lamina propria revealed baseline reductions in neutrophils, a key effector of early bacterial defense, and a failure to expand Th17 cells during infection. Th17 cells are potent inducers of neutrophil recruitment and are critical for bacterial control, particularly in the context of *C. rodentium* [1,2]. Their deficiency in senGC-deficient mice, especially females, was accompanied by a compensatory rise in CD8⁺ T cells, suggesting a shift in immune strategy under conditions of impaired barrier function [20, 21].

Transcriptomic analysis further revealed sex-specific patterns. In females, even at baseline, we observed upregulation of immune and goblet cell–associated gene networks, suggesting a primed mucosal state, although this finding is currently based on a limited number of replicates and requires additional validation.

Upon infection, female *Nlrp6^ΔGC^* mice showed amplified bacterial response pathways, indicative of a compensatory transcriptional program triggered by elevated pathogen burden. In contrast, *Nlrp6^fl/fl^* females exhibited accelerated enrichment of immune-related pathways, consistent with senGC-mediated immune recruitment. Male mice showed enrichment of defense and inflammasome-related networks, which may explain their relatively milder phenotype.

The observation that female mice are more susceptible to intestinal inflammation is consistent with previous reports [22]. Nlrp6-deficient females have previously shown increased sensitivity to DSS-induced colitis [23]. Several factors may contribute to this sex bias. Gut microbiota composition is known to influence susceptibility to *C. rodentium* infection [7], and sex-dependent microbial profiles have been implicated in other disease models [24]. Thus, microbiota-driven differences in immune priming or barrier integrity may underlie the heightened sensitivity observed in females. Hormonal regulation also emerges as another plausible contributor. Estrogen receptor β (ERβ) has been shown to directly upregulate Nlrp6, promoting autophagy and inflammasome assembly in goblet cells and conferring protection against colitis, a mechanism more active in females [25]. Loss of ERβ or Nlrp6 exacerbates colitis in both sexes, but these effects are more pronounced in females which could be linked to mucus deficiencies, mirroring our findings in *Nlrp6^ΔGC^* mice. More broadly, sex hormone receptor signaling is a key regulator of intestinal homeostasis [26], and variability in the estrous cycle may further modulate mucosal responses [27].

The integrity of the mucus barrier is essential for preventing colitogenic inflammation and infection, and its breach occurs in a stepwise fashion. In our previous manuscript, we delineated the sequence of events leading to epithelial colonization, identifying icGCs and the intercrypt mucus they produce as key components of colonization resistance. The intercrypt mucus was the most probable route of *C. rodentium* entry into the epithelium. In the present study, *Nlrp6^ΔGC^* mice exhibited a tendency towards increased mucus penetrability during early stages of infection, suggesting that once the pathogen breaches the intercrypt mucus, it targets crypt spaces for colonization encountering senGCs. Thereby, an absence of senGCs further compromises the exclusionary properties of the inner mucus layer (IML), likely due to the observed reduction in mucin-filled goblet cells in *Nlrp6^ΔGC^* animals which could be potentially driven by the higher crypt associated pathogen burden. Moreover, our data reveal that senGC-deficient mice exhibit diminished baseline mucus secretion during infection, indicating that senGCs could adopt a ‘mucus maintenance’ role under infectious stress, a previously unrecognized and dynamically modulated function that is absent in knockout animals.

Another instance of goblet cell reprogramming uncovered in this study was the infection-induced expansion of *Nlrp6⁺* GCs at the crypt base in wild-type animals. These cells responded to microbial stimuli with enhanced mucus secretion, as evidenced by increased cavitation and ex vivo mucus growth following TLR agonist stimulation. This inducible secretory response was entirely absent in *Nlrp6^ΔGC^*tissue, confirming its dependence on senGCs. We propose that senGCs not only mediate acute microbial sensing and expulsion but also contribute to sustaining mucus barrier integrity during infection through both inducible and baseline secretory programs.

Taken together, our findings position senGCs as a pivotal GC lineage that integrates microbial sensing with mucus barrier regulation. By dissecting their role in both inducible and baseline secretory programs, we reveal senGCs as dynamic responders to infection that not only expel invading pathogens but also drive crypt sterility and epithelial homeostasis. Their absence disrupts immune coordination, accelerates barrier breakdown, and unveils sex-specific vulnerabilities, particularly in females. These insights expand our understanding of GC heterogeneity and highlight senGCs as a critical axis of crypt specific mucosal defense.

## Materials and Methods

### Handling of animal models

All mice used in these experiments were bred in-house under specific pathogen-free conditions and maintained on a 12-hour light/dark cycle with ad libitum access to food and water. Experimental comparisons were conducted exclusively between age- and sex-matched animals. Muc2-iCre mice (unpublished), generated by Lars Vereckee (VIB, Ghent University), were crossed with *Nlrp6^fl/fl^* mice (unpublished), produced by Philip Rosenstiel (Christian-Albrechts-Universität zu Kiel), to generate goblet cell-specific knockouts. Prior to sample collection, animals were anesthetized with isoflurane and euthanized by cervical dislocation. All animal procedures were approved by the Swedish Laboratory Animal Ethical Committee, Gothenburg.

### Preparation of *C. rodentium* inoculum and experimental infection

The streptomycin-resistant *C. rodentium* strain DBS100 was used for all infection experiments to quantify bacterial burden. For colonic whole-mount analyses, mice were infected with *C. rodentium* DBS100-GFP, kindly provided by Bruce A. Vallance, and processed as described in the colonic whole-mount section. Infection inoculum was prepared by culturing bacteria overnight in LB broth at 37°C in a rotating incubator. Mice were orally gavaged with 200 µL of the infection inoculum (1–4 × 10^9^ CFU). At defined time points post-infection, mice were sacrificed, and samples were collected under aseptic conditions to determine *C. rodentium* load across different anatomical sites. Approximately 3 cm of distal colonic tissue was dissected and flushed with 5 mL of sterile PBS. The luminal contents and flushed tissue were collected separately in pre-weighed tubes to calculate weight of the collected tissue or luminal content. All tissue samples were homogenized in sterile PBS using an Ultra-Turrax T10 dispersing instrument (IKA), which was sequentially cleaned between samples using 70% ethanol (x2) followed by sterile PBS (x1). Intestinal contents were completely homogenized in PBS on a Vortex Genie 2 horizontal shaker. Bacterial loads were determined by serial dilution of homogenates and plating on MacConkey agar supplemented with 100 µg/mL streptomycin, followed by overnight incubation at 37°C and quantification of bacterial CFUs. For each sample, a theoretical limit of detection (LOD) was calculated based on the detection of a single colony at the lowest plated dilution. Average LODs for each sample type within an experiment are indicated on all CFU graphs.

### Preparation of tissue for ex vivo analysis

Tissues were processed for ex vivo analysis following previously established protocols [17]. Freshly dissected colonic tissue was flushed with ice-cold, oxygenated Krebs transport buffer (116 mM NaCl, 1.3 mM CaCl₂, 3.6 mM KCl, 1.4 mM KH₂PO₄, 23 mM NaHCO₃, and 1.2 mM MgSO₄; pH 7.4) to remove luminal contents and loose mucus. After excision of the mesenteric tissue, the colon was opened longitudinally, and the muscle layer was carefully separated by microdissection. The prepared tissue was then mounted mucosa-side up in a custom-designed horizontal perfusion chamber, with oxygenated Krebs-glucose buffer (10 mM) in the basolateral compartment and oxygenated Krebs-mannitol buffer (10 mM) in the apical compartment.

### Quantification of IML thickness and penetrability

To evaluate distal colonic inner mucus layer (IML) barrier formation and function, IML thickness and penetrability to fluorescent microbeads were analysed. The tissue was mounted in the perfusion chamber on ice with the mucosal side facing upwards. The tissue was first stained with Rhodamine-conjugated WGA lectin (50 µg/mL; Vector Labs) and 647-conjugated UEA-1 lectin (50 µg/mL; Vector Labs). After a 15-minute incubation on ice, the samples were rinsed with 1 mL of Krebs transport buffer. A suspension of 1-µm diameter Flurosphere crimson microbeads (1:20 dilution; Thermo Fisher) in Krebs transport buffer was then applied to the apical mucosal surface. After allowing the microbeads to settle for 5 minutes, the surface was gently rinsed with 0.5 mL of Krebs transport buffer to remove excess beads. The apical perfusion chamber was subsequently filled with 2 mL of Krebs transport buffer, and the entire setup was transferred to an LSM700 confocal microscope (Carl Zeiss) for imaging.

Confocal z-stacks were collected using a 20× water-immersion objective lens with 405/555 nm laser excitation, with the Zen acquisition software (Carl Zeiss). The resulting image files were imported into Imaris (Bitplane), where fluorescent signals from WGA (marking tissue) and Flurosphere microbeads were used to generate three-dimensional isosurfaces. The z-axis positions of the tissue surface and individual microbeads were extracted for analysis. IML thickness was determined by calculating the mean distance along the z-axis between the tissue surface and the microbeads. To assess mucus penetrability, the spatial distribution of microbeads within the mucus layer was analysed. For each z-stack, a frequency distribution of microbead distances relative to the tissue surface was generated using R. To enable comparisons between samples, each curve was normalized to its maximum frequency value to correct for differences in total bead counts per z-stack. Curves were also normalized to the position of the mucus surface (calculated from the z-position with the highest microbead density) to account for variations in mucus thickness, excluding beads located above the mucus layer from analysis. Finally, the area under each normalized curve was calculated to quantify microbead penetrability into the mucus layer across samples.

### Ex vivo quantification of IML secretion and growth rate

The secretion and processing of the inner mucus layer (IML) in the distal colon were assessed by measuring the rate of mucus growth ex vivo, as described previously [17]. Tissue samples were mounted in custom perfusion chambers with the basolateral compartment continuously perfused at a rate of 5 mL/h, and the system was maintained at 37°C. To visualize mucus formation on the apical surface, 10µm black latex microbeads (Polybead; diluted 1:20) were applied on top of the tissue. Mucus thickness was determined using a stereomicroscope by measuring the distance between the epithelial surface and the microbeads with a glass micropipette connected to a micrometer. Measurements were recorded at 15-minute intervals over a 60-minute period (t = 0 to 60 min). At *t* = 30 minutes, tissues were apically treated with the TLR2 agonist P3CSK4 (50 µg/ml) or LPS (ultrapure, 200 µg/ml, Invivogen) or Carbachol (1mM, Invivogen). Mucus growth was quantified at *t* = 45 minutes (15 minutes post-treatment) to assess the impact of stimulation on secretion rate. All microbial-associated molecular patterns (MAMPs) were reconstituted in double-distilled water (ddH₂O), sonicated, and stored at −20 °C until use. For experiments involving pharmacological inhibition of MAMP-induced mucus secretion, inhibitory compounds were added to the apical buffer at *t* = 0 and maintained throughout the assay. Dynasore (100 µM; Merck) was used to block endocytosis-dependent signaling pathways. Following fixation in 4% formalin overnight and transfer to 70% ethanol, tissues were processed for paraffin embedding and sectioning. Sections were subsequently stained using Alcian Blue/Periodic Acid–Schiff (AB/PAS) to visualize mucin-filled goblet cells.

### Ex vivo analysis of Fecal Mucus Layer (FML)

Stool samples from *C. rodentium*-infected and control mice were collected during colon dissection and immediately placed in Krebs transport buffer. To stabilize the stool during imaging, a custom slide was prepared by shaping tack clay into a dome with a central indentation. A stool pellet was carefully positioned within the indentation, and a chamber was placed over it such that the apical portion of the stool protruded slightly outside the chamber. The mucus layer encapsulating the stool was stained using a solution containing Rhodamine-conjugated WGA lectin (50 µg/mL; Vector Labs) and 647-conjugated UEA-1 lectin (50 µg/mL; Vector Labs). After a 15-minute incubation, the chamber was gently rinsed with 0.5 mL of Krebs transport buffer to remove excess stain. The apical compartment of the chamber was then filled with 2 mL of Krebs buffer, and the setup was transferred to an LSM700 confocal microscope (Carl Zeiss) for imaging. Confocal z-stacks of the fecal mucus layer were acquired using a 20× water-immersion objective lens with 555 nm and 647 nm laser excitation, controlled via Zen acquisition software (Carl Zeiss). The images were analysed using Imaris (Bitplane), where the mucus surface was mapped based on the WGA signals. Measurements of mucus volume, length, and width were obtained, and the thickness (height) of the mucus layer surrounding the stool was calculated using the formula:

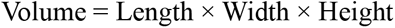

### Colonic whole-mount preparation and imaging

Colonic tissue was harvested from mice infected with *C. rodentium* GFP-DBS100 or from uninfected ROSA-tdRFP/Muc2-iCre reporter mice. Tissue collection, fixation and processing were performed identically for both experimental groups. Collected samples were immediately placed in oxygenated Krebs transport buffer (pH 7.2) on ice until dissection. For dissection, tissues were transferred to a dissection tray and flushed thoroughly with Krebs buffer to remove luminal contents and loose mucus. The mesentery was removed, and the colon was opened longitudinally. The Krebs buffer was removed, and the tissue was fixed by immersion in 4% neutral buffered formalin for 1 hour at room temperature. Following fixation, samples were rinsed with PBS and permeabilized in 0.5% Triton X-100 in PBS for 15 minutes at room temperature with gentle shaking. The permeabilization solution was then removed, and tissues were rinsed again with PBS. Staining was performed as necessary by incubating permeabilized tissues for 1 hour at room temperature with gentle shaking in a solution containing 1:500 Hoechst-34580 (Thermo Fisher Scientific), 50 µg/mL UEA1 (Rhodamine), and 50 µg/mL WGA (Alexa647) in PBS. Where indicated in experimental protocols, Phalloidin (Invitrogen, A22287; 1:1000 dilution) was used as a marker to visualize the tissue surface. Detection of *C. rodentium* was performed using an anti-GFP antibody (Mouse, A-11122; 1:200) when required. After staining, tissues were washed twice in PBS for 15 minutes each and mounted on glass slides with the mucosal surface facing upwards. Confocal imaging was carried out using an LSM700 microscope (Zeiss).

### Colonic swiss roll preparation and quantification of goblet cell number and crypt length

Mice infected with *C. rodentium* DBS100-GFP were sacrificed, and the entire colon was collected in oxygenated Krebs buffer. The mesentery was removed, and the colon was thoroughly cleaned to remove luminal contents. The colon was then cut open longitudinally and rolled into a Swiss roll from distal to proximal, with the mucus layer facing upward. Rolls were fixed in 4% neutral buffered formalin for 24 hours, transferred to 70% ethanol, and sent for paraffin embedding and sectioning (Histocenter AB). Histological staining with Alcian Blue–Periodic Acid-Schiff (AB-PAS) was performed by Histocenter. Slides were scanned using a NanoZoomer-SQ Digital Slide Scanner C13140-01 (Hamamatsu), and images were analysed with ImageJ. Ten biological replicates, imaged at different time points (uninfected, and 3-, 7-, and 10-days post-infection), were included in the analysis. Each Swiss roll was divided into four regions: proximal colon (striated colonic region), proximal mid-colon, mid-colon, and distal colon. Each region was analysed separately. For each region, the lengths of 15 crypts were measured, and goblet cell numbers were counted from 10 individual crypts. The average crypt length and goblet cell count were then calculated for each animal across the four regions. Goblet cell (GC) frequency was expressed as average GC count per crypt divided by average crypt length.

### RNAscope and immunohistochemical staining

For RNA *in situ* hybridization, formalin-fixed Swiss roll tissue sections were processed using the RNAscope technique (Advanced Cell Diagnostics) according to ACD Biotech’s instructions [5]. Following hybridization, sections were washed three times with TBS-Tween20 and blocked with 1% BSA in Tween-20 for 20 minutes at room temperature. Sections were then incubated overnight at 4°C with stated primary antibodies: Anti-EpCAM (Rabbit, HPA026761-25UL; 1:200) and Anti-GFP (Mouse, A-11122; 1:200) to label the respective targets. After washing with TBS-Tween20, primary antibodies were detected using goat anti-rabbit Alexa Fluor 488 or Alexa Fluor 555–conjugated secondary antibodies (1:2000; Thermo Fisher) for 2 hours at room temperature. Finally, slides were rinsed with double-distilled water and counterstained with Hoechst dye (5 µg/mL; Sigma Aldrich) for 5 minutes. Stained sections were imaged using an LSM700 confocal microscope (Zeiss).

### Isolation of lamina propria (LP) cells from mouse distal colon

A 5–6 cm segment of the distal large intestine, encompassing the mid and distal colon, was dissected. The segment was flushed using a syringe fitted with a blunt needle containing cold 10% v/v 10× Hank’s Balanced Salt Solution without Mg²⁺ and Ca²⁺, supplemented with 1 mM HEPES (HBSSwo), to remove luminal contents. During dissection and transport, distal colon samples were kept in HBSSwo on ice. Mesenteric fat and blood vessels were carefully removed. To prepare for digestion, the tissue was turned inside out using negative pressure from a syringe with a blunt needle to expose the luminal surface. Both ends of the inverted colon were ligated with common thread: one end was tied first, the lumen inflated with HBSSwo, and the opposite end tied off. This ensured full opening of crypts, providing optimal access of digestive solutions to all epithelial cells, including those at the crypt base. For digestion, the inflated colon was incubated in 20 mL of pre-warmed HBSSwo containing 5 mM EDTA, 5% v/v heat-inactivated fetal bovine serum, and 1 mM freshly prepared dithiothreitol (DTT; Thermo Fisher Scientific, R0861) at 37°C with shaking at 140 rpm for 30 minutes. Following this, epithelial cells were detached by vigorous shaking for 30 seconds. The digestion uspernatent was stored at 4°C until the completion of the next step. The remaining tissue was then transferred to fresh pre-warmed pre-digestion buffer and incubated under the same conditions for an additional 30 minutes. The supernatants from both digestions were pooled and stored at 4°C. Residual tissue was discarded. The pooled cell suspension (∼40 mL) was centrifuged at 400 × g for 10 minutes at 4°C, and the supernatant discarded. The resulting pellet was resuspended in 1 mL of Collagenase Type I (2 mg/mL; Sigma Aldrich, 637958) with 4 μL of DNase I (200 U/mL; Zymo Research, E1009) and incubated at room temperature for 30 minutes, with gentle pipetting every 5 minutes. Following enzymatic digestion, cells were washed with 25 mL of FACS buffer (HBSSwo supplemented with 2% FBS and 1 mM EDTA), centrifuged again at 400 rcf for 10 minutes at 4°C. The supernatant was discarded and the pellet resuspended in the desired volume of FACS buffer for downstream applications.

To continue with LP immune cell preparation, the remaining intestinal tissue was washed once in PBS, cut into small pieces, and resuspended in 40 mL of R5 medium containing Dispase (160 μL of 50 U/mL stock) and Type VIII Collagenase (100 μL of 10,000 U/mL stock prepared as 200 mg in 7.6 mL TC-PBS) per intestine. Samples were incubated for 1.5–2 hours (not less than 1 h 30 min and not more than 2 h 15 min) at 37 °C on a rotator at 140 rpm, with vigorous hand-shaking every 20–30 minutes. Following digestion, tissue suspensions were passed through a 70 μm cell strainer and centrifuged at 4000 rpm for 5 minutes. Cell pellets were pooled, resuspended in 40 mL of PBS and stored on ice until further staining and downstream applications.

### Staining of lamina propria cells for flow cytometry

Cell viability was assessed using eFluor® 780 (AH Diagnostics, 65-0865-14), diluted 1:1000 in PBS, added to the cells and incubated for 30 minutes on ice in the dark. After incubation, cells in the 96-well plate were centrifuged at 1500 rpm for 5 minutes at 4°C, and the supernatant was discarded. Depending on the target molecule and antibody type, fixation was performed as required. For extracellular markers, staining was done prior to fixation, either directly or indirectly, depending on whether the antibody was pre-conjugated. Antibodies were diluted in 200 µL of FACS buffer at optimized concentrations (see below for a list of antibodies and lectins used in the study). Staining was carried out for 30 minutes on ice in the dark. Flow cytometry was performed using a BD FACS Symphony A1, and data were subsequently analysed with FlowJo software. The antibodies used for flow cytometry included Alexa Fluor 700-conjugated anti-CD45 (1:1000; BioLegend, 304023), BUV395-conjugated anti-Siglec-F (1:1000; BD OptiBuild, BDB740280), PerCP-eFluor 710-conjugated anti-Ly6G (1:1500; eBioscience, 46-9668-82), APC-conjugated anti-CD3 (1:1000; eBioscience, 17-0032-82), Brilliant Violet 510-conjugated anti-CD4 (1:1000; BioLegend, 100449), PE-conjugated anti-CD8α (1:1000; BioLegend, 100708), PE/Cyanine5-conjugated anti-CD335 (NKp46; 1:1250; BioLegend, clone 29A1.4), and Brilliant Violet 605-conjugated anti-CD11b (1:1000; BioLegend, clone M1/70).

Additional antibodies included APC-conjugated anti-Ly6C (1:1000; BioLegend, clone HK1.4), Brilliant Violet 421-conjugated anti-F4/80 (1:750; BioLegend, clone BM8), Brilliant Violet 711-conjugated anti-CD64 (1:1000; BioLegend, clone X54-5/7.1), APC-eFluor 780-conjugated anti-MHCII (1:1000; eBioscience, 47-5321-82), PE-conjugated anti-CD11c (1:1000; BioLegend, clone N418), Brilliant Violet 650-conjugated anti-XCR1 (1:1500; BioLegend, 148220), PE/Cyanine7-conjugated anti-SIRPα (CD172a; 1:1500; BioLegend, 144007), FITC-conjugated anti-CD19 (1:1000; BioLegend, 115505), and PE-conjugated anti-CD138 (Syndecan-1; 1:1500; BioLegend, 142503). For intracellular transcription factor staining, Alexa Fluor 488-conjugated anti-FoxP3 (1:1000; BioLegend, 126405), PE-conjugated anti-GATA3 (1:1000; BioLegend, 653803), and PE/Cyanine7-conjugated anti-T-bet (1:1000; BioLegend, 644823) were used

### DSS-induced colitis

Experimental colitis was induced in Nlrp6^fl/fl^ and Nlrp6^ΔGC^ mice by providing 3% (wt/vol) dextran sulfate sodium (DSS; TdB Labs) in drinking water ad libitum for 8 days. Body weight was recorded daily throughout treatment to ensure mice did not lose more than 15% of their initial body weight. At the experimental endpoint, colon length and spleen weight were measured. Colonic tissues were flushed, opened longitudinally and prepared as Swiss rolls, followed by formalin fixation, paraffin embedding, sectioning and hematoxylin and eosin (H&E) staining. Histopathological changes in the mid-distal colon were evaluated using a modified scoring system [28], assessing inflammatory infiltration, GC depletion, crypt hyperplasia, muscular thickening, submucosal inflammation and ulceration. Histological assessments were performed by blinded analysis, and the averaged scores for each parameter were combined to obtain an overall histopathology score.

### RNA Extraction, Bulk RNA Sequencing, and Analysis

Distal colon samples from *Nlrp6^fl/fl^* and *Nlrp6^ΔGC^*mice infected with WT *C. rodentium* were collected, immediately placed in RNAlater (Qiagen), and stored at −20°C until processing. For RNA extraction (RNeasy Plus Mini Kit, Qiagen), tissues were thawed, minced, and homogenized in RLT Plus buffer containing β-mercaptoethanol using a rotor–stator. Lysates were briefly clarified by centrifugation and passed through a gDNA Eliminator spin column to remove genomic DNA. The flow-through was mixed 1:1 with 70% ethanol and loaded onto RNeasy spin columns, followed by sequential washes with Buffer RW1 and Buffer RPE (including the second RPE wash and an additional dry spin to remove residual ethanol). Where indicated, an on-column RNase-free DNase treatment was included prior to RW1. RNA was eluted in 20 µL of RNase-free water, quantified, and purity assessed using NanoDrop. Samples were sequenced by Novogene. Sequencing reads were quality-checked and trimmed to remove adapters and low-quality bases, then aligned and quantified against the GRCm39 *mus musculus* reference genome to obtain gene-level counts. Lowly expressed genes were filtered, and samples were normalized. Principal component analysis (PCA) on variance-stabilized data was used to assess baseline sex effects and infection trajectories. Differential expression was tested for using DESeq2 using a design accounting for sex and genotype with two analyses [29]:

1. Baseline (uninfected) comparison of *Nlrp6^ΔGC^* vs *Nlrp6^fl/fl^* for each sex.
2. Infection response within each genotype by contrasting infected vs genotype-matched uninfected samples at 3, 7, and 10 dpi.

DEGs were defined at a Benjamini–Hochberg FDR adjusted p-value (padj) <0.05. For uniqueness analyses to identify genotype-specific versus shared DEG sets at each time point, DEGs were further filtered by log₂ fold-change >1. Functional analysis of different DEG sets was performed using the PANTHER Classification System overrepresentation test (GO Biological Process Complete, Fisher’s Exact test, multiple testing: Benjamini–Hochberg FDR).

### RNA extraction and qPCR

Colonic tissue samples were collected in RNAlater (Qiagen) and stored until RNA extraction. Tissues were transferred into 350 µL of RLT buffer (Qiagen) containing β-mercaptoethanol (Gibco) and homogenized using an Ultra-Turrax T10 dispersing instrument (IKA). Total RNA was isolated using RNeasy Plus Mini columns (Qiagen) according to the manufacturer’s instructions and eluted in 20 µL of RNase-free water.

RNA concentration and purity were assessed using a NanoDrop 2000/2000c spectrophotometer (Thermo Scientific), and samples were stored at −80°C until further analysis.

Complementary DNA (cDNA) was synthesized from 600 ng of RNA using the High-Capacity cDNA Reverse Transcription Kit (Thermo Fisher Scientific). Quantitative real-time PCR (qRT-PCR) was performed using a CFX96 Real-Time PCR Detection System (Bio-Rad), with an initial denaturation at 95°C for 2 minutes, followed by 40 cycles of 95°C for 5 seconds and 58°C for 30 seconds. Data were analysed using CFX Manager software (version 3.1; Bio-Rad). Relative *Nlrp6* expression was calculated using CFX Manager software (version 3.1; Bio-Rad), with normalization to the reference genes Gapdh and Rplp0. *Nlrp6* expression was assessed using a prevalidated mouse primer assay (Bio-Rad; qMmuCID0024291). The primer sequences (5′-3′) were *Gapdh*: forward, GGAGAAACCTGCCAAGTATG and reverse, GGAGTTGCTGTTGAAGTCG; *Rplp0*: forward, GCGACCTGGAAGTCCAACTA and reverse, TCTCCAGAGCTGGGTTGTTT.

### Analysis of scRNA -seq

A previously published single-cell RNA-sequencing dataset (GEO accession: GSE157313) was reanalyzed using Loupe Browser (version 6; 10X Genomics) [8]. Cells were filtered to exclude those with mitochondrial gene content above 20%, fewer than 200 or more than 5,500 detected genes, and those with over 6,500 unique gene counts in the distal colon. Immune and enteroendocrine cells were removed based on the expression of Ptprc and Chgb, respectively. To reduce contamination from enterocytes and colonocytes, only cells with Muc2 expression levels above 2 were retained. Following these quality control steps, 15,392 genes across 6,123 distal colon cells were included in the final analysis. Dimensionality reduction was performed using Uniform Manifold Approximation and Projection (UMAP), followed by k-means clustering (K = 10), which identified five distinct goblet cell clusters. Cluster identities were assigned based on previously published annotations [8].

### Statistical analyses

Details of the statistical analyses performed for each experiment are provided in the corresponding figure legends. The term “n” refers to the number of biological replicates (individual animals) used in each experiment. Transcriptomic visualizations were generated using Loupe Browser version 6. Statistical analyses and data visualization were performed using GraphPad Prism version 9.4.1 (GraphPad Software). Results are presented as either mean ± standard deviation (SD) or median with range, as specified. The statistical tests used for each analysis are indicated where applicable. Significance levels are denoted as follows: * P < 0.05; ** P < 0.01; *** P < 0.001; **** P < 0.0001; ns or “not specified” indicates not significant.

### Software used

The software used for experimental data acquisition, processing, visualization and analysis included ZEN (version 2.3; Carl Zeiss, www.zeiss.com), Imaris x64 (version 9.5.0; Oxford Instruments, imaris.oxinst.com), GraphPad Prism (version 9.4.1; GraphPad Software, www.graphpad.com), Loupe Browser (version 6; 10x Genomics, www.10xgenomics.com), ImageJ (Fiji; fiji.sc) and FlowJo (FlowJo LLC, www.flowjo.com).

